# Human B cell clonal expansion and convergent antibody responses to SARS-CoV-2

**DOI:** 10.1101/2020.07.08.194456

**Authors:** Sandra C. A. Nielsen, Fan Yang, Katherine J. L. Jackson, Ramona A. Hoh, Katharina Röltgen, Bryan Stevens, Ji-Yeun Lee, Arjun Rustagi, Angela J. Rogers, Abigail E. Powell, Javaria Najeeb, Ana R. Otrelo-Cardoso, Kathryn E. Yost, Bence Daniel, Howard Y. Chang, Ansuman T. Satpathy, Theodore S. Jardetzky, Peter S. Kim, Taia T. Wang, Benjamin A. Pinsky, Catherine A. Blish, Scott D. Boyd

**Author notes:** These authors contributed equally (Sandra C. A. Nielsen, Fan Yang, Katherine J. L. Jackson, Ramona A. Hoh). Correspondence (C.A.B.), (S.D.B.).

## Abstract

During virus infection B cells are critical for the production of antibodies and protective immunity. Here we show that the human B cell compartment in patients with diagnostically confirmed SARS-CoV-2 and clinical COVID-19 is rapidly altered with the early recruitment of B cells expressing a limited subset of IGHV genes, progressing to a highly polyclonal response of B cells with broader IGHV gene usage and extensive class switching to IgG and IgA subclasses with limited somatic hypermutation in the initial weeks of infection. We identify extensive convergence of antibody sequences across SARS-CoV-2 patients, highlighting stereotyped naïve responses to this virus. Notably, sequence-based detection in COVID-19 patients of convergent B cell clonotypes previously reported in SARS-CoV infection predicts the presence of SARS-CoV/SARS-CoV-2 cross-reactive antibody titers specific for the receptor-binding domain. These findings offer molecular insights into shared features of human B cell responses to SARS-CoV-2 and other zoonotic spillover coronaviruses.

## INTRODUCTION

The novel human severe acute respiratory syndrome coronavirus 2 (SARS-CoV-2) is the etiological agent of the coronavirus disease 2019 (COVID-19) (Huang et al., 2020; Zhu et al., 2020) pandemic. Prior to the emergence of SARS-CoV-2, six human coronaviruses (hCoVs) were known; four seasonal hCoVs (hCoV-229E, -NL63, -HKU1, and -OC43) (Su et al., 2016) causing usually mild upper respiratory illness, and the two more recently discovered SARS-CoV (Peiris et al., 2003) and MERS-CoV (Zaki et al., 2012) viruses that arose from spillover events of virus from animals into humans. It is expected that humans are naïve to SARS-CoV-2 and will display a primary immune response to infection. Humoral immune responses will likely be critical for the development of protective immunity to SARS-CoV-2. Recently, many novel SARS-CoV-2 neutralizing antibodies from convalescent COVID-19 patients have been reported (Cao et al., 2020; Ju et al., 2020; Robbiani et al., 2020b; Wu et al., 2020b), which offer an important resource to identify potential protective or therapeutic antibodies. However, a deeper understanding of the B cell antigen receptors that are stimulated and specific to this acute infection is needed to define the shared or distinct features of humoral responses elicited compared to other viral infections, and to assess the extent to which responses to SARS-CoV-2 have breadth extending to other coronaviruses within the subgenus Sarbecovirus.

## RESULTS

### SARS-CoV-2 infection causes global changes in the antibody repertoire

High-throughput DNA sequencing of B cell receptor heavy chain genes defines clonal B cell lineages based on their unique receptor sequences, and captures the hallmarks of clonal evolution, such as somatic hypermutation (SHM) and class switch recombination during the evolving humoral response (Zhou and Kleinstein, 2019). To study the development of SARS-CoV-2-specific humoral responses, we collected a total of 38 longitudinal peripheral blood specimens from 13 patients, sampled at a median of 3 time points (range 1-5) admitted to Stanford Hospital with COVID-19 confirmed by quantitative reverse transcription PCR (RT-qPCR) testing. The times of blood sampling were measured as days post symptom onset (DPSO). All patients exhibited SARS-CoV-2 receptor-binding domain (RBD)-specific IgA, IgG, and IgM antibodies (Table 1). Immunoglobulin heavy chain (IGH) repertoires were sequenced and compared to a healthy human control (HHC) data set from 114 individuals (Nielsen et al., 2019). An example of data from a HHC individual matched by mean sequencing depth of reads and B cell clones across the COVID-19 cohort is shown in Figure 1 (top panel). In healthy subjects at baseline, IgM and IgD sequences are primarily derived from naïve B cells with unmutated IGHV genes, whereas class switched cells expressing IgA or IgG subtypes have elevated SHM. In contrast, SARS-CoV-2 seroconverted patients (blue labels in Figure 1), show a highly polyclonal burst of B cell clones expressing IgG, and to a lesser extent IgA, with little to no SHM. Longitudinal data from a patient prior to and after seroconversion shows an increase in the proportion of class switched low SHM clones (bottom panels in Figure 1). Seronegative samples (red labels in Figure 1) show IGH repertoires similar to uninfected HHC, suggesting an earlier stage in the infection for these particular patients at these time points.

**Table 1.** Individual COVID-19 patient sample information in days post symptom onset (DPSO). Seroconversion was determined by ELISA to the SARS-CoV-2 RBD antigen (STAR METHODS).

| Patient ID | DPSO | IgA seroconversion | IgG seroconversion | IgM seroconversion | gDNA library | cDNA library |
| --- | --- | --- | --- | --- | --- | --- |
| 7450 | 9 | Yes | Yes | Yes | Yes | Yes |
| 7450 | 22 | Yes | Yes | Yes | Yes | Yes |
| 7450 | 25 | Yes | Yes | Yes | Yes | No |
| 7450 | 27 | Yes | Yes | Yes | Yes | Yes |
| 7451 | 8 | No | No | No | Yes | Yes |
| 7452 | 18 | Yes | Yes | Yes | Yes | Yes |
| 7453 | 8 | No | No | No | Yes | Yes |
| 7453 | 11 | No | No | No | Yes | Yes |
| 7453 | 15 | Yes | Yes | Yes | Yes | Yes |
| 7453 | 18 | Yes | Yes | Yes | Yes | Yes |
| 7453 | 20 | Yes | Yes | Yes | Yes | No |
| 7454 | 16 | Yes | Yes | Yes | Yes | Yes |
| 7455 | 9 | Yes | Yes | Yes | Yes | Yes |
| 7455 | 11 | Yes | Yes | Yes | Yes | Yes |
| 7455 | 12 | Yes | Yes | Yes | Yes | Yes |
| 7455 | 16 | Yes | Yes | Yes | Yes | Yes |
| 7455 | 19 | Yes | Yes | Yes | Yes | Yes |
| 7480 | 11 | Yes | Yes | Yes | Yes | Yes |
| 7480 | 14 | Yes | Yes | Yes | Yes | Yes |
| 7480 | 17 | Yes | Yes | Yes | Yes | No |
| 7481 | 32 | Yes | Yes | Yes | Yes | Yes |
| 7481 | 35 | Yes | Yes | Yes | Yes | No |
| 7481 | 37 | Yes | Yes | Yes | Yes | No |
| 7482 | 5 | No | No | No | Yes | Yes |
| 7482 | 8 | No | No | No | Yes | Yes |
| 7482 | 10 | No | No | Yes | Yes | Yes |
| 7482 | 12 | No | Yes | Yes | Yes | Yes |
| 7483 | 35 | Yes | Yes | Yes | Yes | Yes |
| 7483 | 40 | Yes | Yes | Yes | Yes | Yes |
| 7484 | 11 | Yes | Yes | Yes | Yes | Yes |
| 7484 | 14 | Yes | Yes | Yes | Yes | Yes |
| 7484 | 17 | Yes | Yes | Yes | Yes | Yes |
| 7485 | 8 | No | Yes | Yes | No | No |
| 7485 | 10 | Yes | Yes | Yes | Yes | Yes |
| 7485 | 12 | Yes | Yes | Yes | Yes | Yes |
| 7486 | 14 | No | Yes | Yes | Yes | Yes |
| 7486 | 17 | Yes | Yes | Yes | Yes | Yes |
| 7486 | 19 | Yes | Yes | Yes | Yes | Yes |

**Figure 1.**
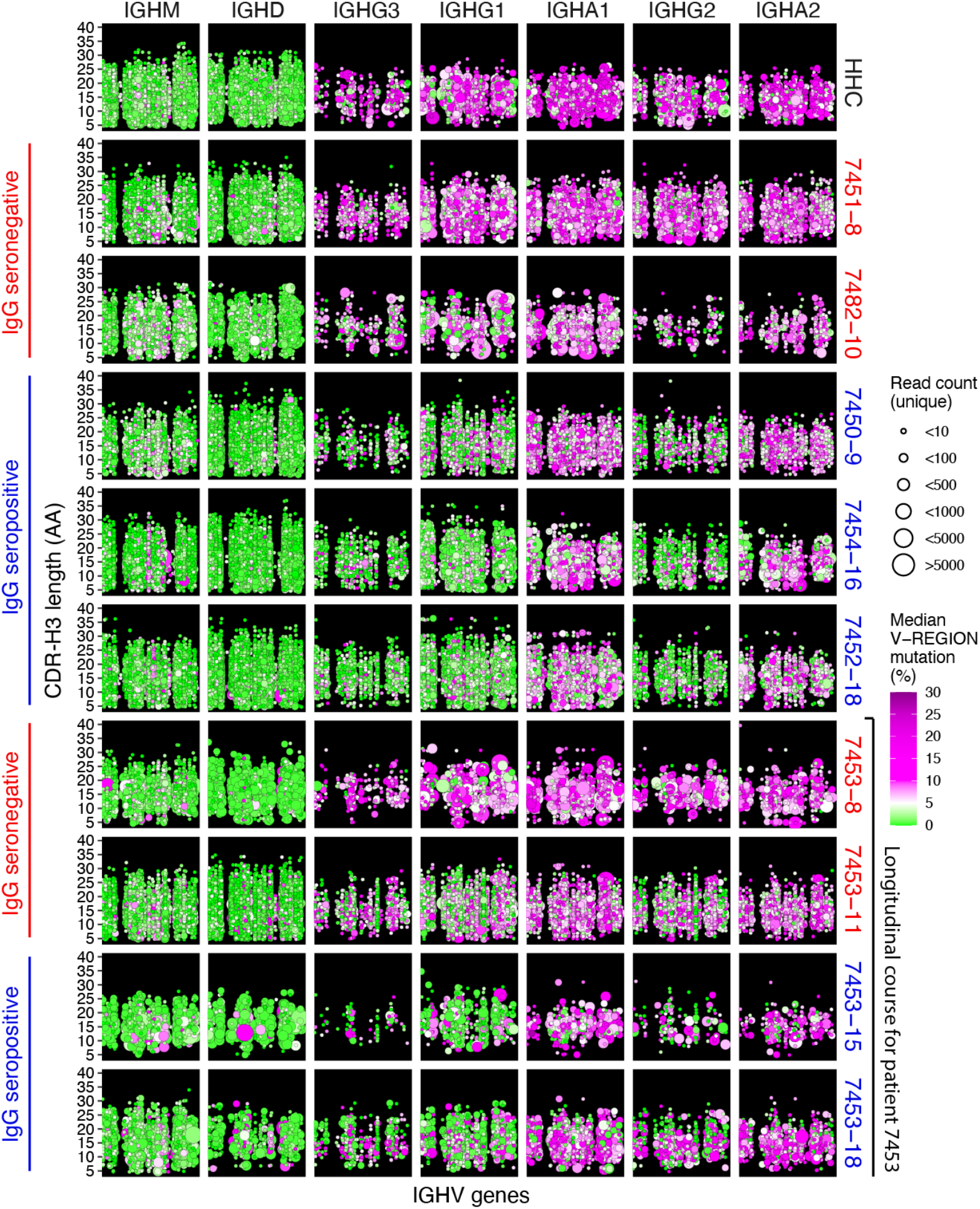
COVID-19 patient IGH repertoires show early and extensive class-switching to IgG and IgA subclasses without significant somatic mutation. Points indicate B cell clonal lineages, with the position denoting the clone’s isotype (panel column), human healthy control (HHC) or patient ID (panel row), IGHV gene (x-axis, with IGHV gene in the same order and position in panels, but not listed by name due to space constraints), and CDR-H3 length in amino acids (AA) (y-axis within each panel). The point color indicates the median IGHV SHM frequency for each clone and the size indicates the number of unique reads grouped into the clone. Points are jittered to decrease over-plotting of clones with same IGHV gene and CDR-H3 length. Patient label colors indicate sample IgG seroconversion (blue) or seronegative (red) for the displayed sample with the number following the patient ID corresponding to days post symptom onset. The final four rows of panels show the IGH repertoire changes within a single participant (7453) prior to and after seroconversion.

The increased fraction of unmutated and low mutation (<1% SHM in IGHV gene) clones among the class switched IgG subclasses in seroconverted COVID-19 patients compared to HHC was statistically significant (IgG1: p-value = 1.884e-08; IgG2: 1.554e-08; IgG3: 3.754e-08; IgG4: 0.00044) (Figures 2A and 2B). The detailed SHM frequencies and fractions of unmutated clones of each sample are shown in Figure S1. While most B cells in COVID-19 patients prior to seroconversion showed IgG SHM frequencies comparable to HHC, there is a rapid increase in the proportion of IgG-expressing (IgG+) B cells with low SHM during the DPSO (Figure 2A, lower panels). Notably, prior to seroconversion, B cells expressing a few IGHV genes, particularly IGHV3-30-3 and IGHV1-2, showed earlier changes than the rest of the repertoire, with increased IgG class switched low-SHM clones (Figures 2C and 2D). IGHV3-9 showed a similar trend but was not significant (Figures 2C and 2D).

**Figure 2.**
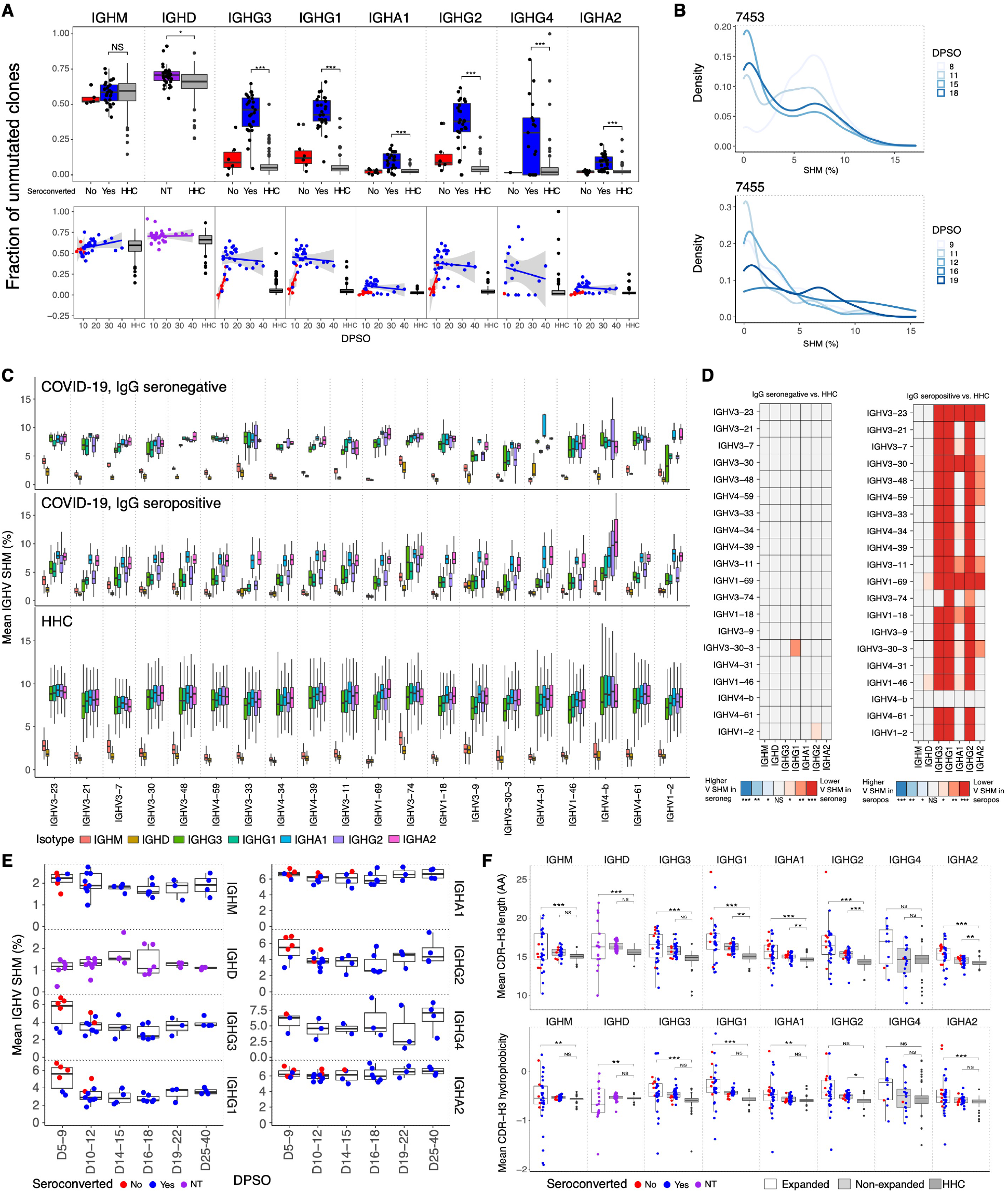
IGH repertoire signatures of SARS-CoV-2 infection. (A) Fraction of unmutated (<1% SHM) B cell lineages for each isotype subclass grouped by seroconversion status (top panel) or plotted by days post symptom onset (DPSO, bottom panel). Colors indicate patient sample serology: not tested (NT, purple), seronegative (red), and seropositive (blue) and are plotted specific to the isotype tested. Points are shown for all COVID-19 samples, whereas only outliers are displayed for the 114 healthy human controls (HHCs). Differences between the seropositive group and HHC was tested using two-sided Wilcoxon–Mann-Whitney (for patients with more than one sample, the mean value of these was used). (B) Distribution of clone percentage SHM plotted as kernel density for clones detected at multiple time points from patients 7453 and 7455. Lines are colored by DPSO. (C) Mean IGHV SHM percent for each isotype subclass observed for the IgG seronegative patient samples (top), IgG seropositive samples (middle) or HHCs (bottom). IGHV order is based on the 20 most common IGHV genes in IgM in the patients and isotypes are plotted by their chromosomal ordering. The plot axes were chosen to show the box-whiskers on a readable scale; rare outlier points with extreme values are not shown but were included in all analyses. (D) Heatmap of patient IGHV gene SHM for seroconverted and non-seroconverted samples compared to HHC using paired Wilcoxon tests with Bonferroni correction for multiple hypothesis testing. The color scale encodes the significance level and whether the SHM was higher (blues) or lower (reds) in COVID-19 relative to HHC. (E) Longitudinal SHM for each isotype subclass for COVID-19 patients are plotted binned by DPSO. Points are colored for each sample’s seroconversion status and boxplots summarize median and interquartile ranges. (F) Mean CDR-H3 length (top panel) and mean CDR-H3 hydrophobicity (bottom panel), COVID-19 patient samples grouped by expanded clones (white) or non-expanded clones (light grey), and total clones from HHC (dark grey). Differences between the expanded/non-expanded groups and HHC were tested using one-way ANOVA with Tukey’s HSD test. (A), (D), and (F) ***p-value < 0.001; **p-value < 0.01; *p-value ≤ 0.05; NS: p-value > 0.05.

We previously observed a similar influx of low mutation clones into the IgG compartment in acute Ebola virus (EBOV) infection (Davis et al., 2019), but in EBOV acute viral infection there was a prolonged delay (lasting months) in accumulation of SHM in those clones. In contrast, examination of clones detected at two or more time points show that after the initial appearance of low-SHM clones, the COVID-19 patients show increases in the proportion of IgG+ B cells with intermediate SHM frequencies (2-5%) within the first three weeks post-onset of symptoms in the patients for whom the longest time courses were observed (patients 7453 and 7455, Figure 2B). Similarly, examination of the total clones within each isotype shows the appearance of low-SHM clones post-seroconversion within the first two weeks post-onset of symptoms, and subsequent increases in SHM over the following two weeks (Figure 2E). In further contrast to EBOV, COVID-19 primary infection stimulated polyclonal B cell responses with both IgG and, in some patients, IgA subclasses, rather than IgG alone (Figure 1). Overall, among IgG+ B cells in COVID-19 patients, the proportion of IgG1+ cells was increased, with decreases in IgG2 and IgG3, and median usage of IgG1 was 1.7-fold greater than that seen in HHC B cells (Table S1).

Comparison of the IGHV genes used by COVID-19 and HHC individuals (Figure S2) revealed skewing of the responding IGH repertoires away from frequently utilized IGHV genes in HHC, such as IGHV3-7, IGHV3-23, and IGHV5-51, and enrichment of IGHV1-24, IGHV3-9, IGHV3-13, and IGHV3-20 in IgG+ B cells. The preferential selection of B cells using particular IGHV genes has been observed in other antiviral responses, such as the preference for IGHV1-69 in response to some influenza virus antigens (Avnir et al., 2016). Highly utilized IGHV genes in IgG-seroconverted COVID-19 patients display low median SHM in IgG1-switched B cells (range 2.4-7.5%), compared to higher median IgA1 SHM (range 5.8-9.5%) (Figure 2C).

In the SARS-CoV-2 stimulated B cell proliferations, high-frequency expanded clones detected in two or more replicate IGH sequence libraries generated from separate aliquots of template showed increased proportions of low-SHM members (Figures S3A and S3B, IgG1: p-value = 0.005, IgG2: 0.014, IgG3: 0.036). Expanded clones also had longer and more hydrophobic IGH complementarity-determining region-3 (CDR-H3) sequences in the class switched isotypes compared to HHC (Figure 2F), highlighting IGHV features selected in B cells responding to SARS-Cov-2 infection and consistent with a rapid proliferation of cells recently differentiated from naïve B cells (Grimsholm et al., 2020). CDR-H3 charge and aromaticity showed modest differences in COVID-19 patients compared to HHC, including more negative charge in non-expanded clones (Figure S3C). Notably, the relative IGHV gene usage frequencies in expanded clones compared to non-expanded clones of COVID-19 patients showed a different pattern than overall IGHV gene usage, with IGHV1-24, IGHV3-13, and IGHV3-20 frequencies that were increased in the total repertoire but used less often in expanded clones, suggesting that the B cells expressing these IGHV genes are highly polyclonal with small clone sizes. Eleven IGHV genes were significantly enriched in expanded clones versus non-expanded clones (Figure S3D) suggesting preferential recruitment of B cells and viral epitope binding by IGH using these germline IGHV segments.

### Convergent antibody rearrangements are elicited in COVID-19 patients

Despite the diversity of antigen-driven antibody responses, we and others have previously identified patterns of highly similar, “convergent” antibodies shared by different individuals in response to pathogens such as EBOV (Davis et al., 2019) or Dengue virus (Parameswaran et al., 2013). Such convergent antibodies make up a small proportion of the total virus-specific B cell response in each individual (Davis et al., 2019). To identify putative SARS-CoV-2-specific antibody signatures, we analyzed clones with shared IGHV, IGHJ, and CDR-H3 region length, and clustered the CDR-H3 sequences at 85% amino acid identity using CD-HIT (Fu et al., 2012), to find clusters spanning two or more COVID-19 patients and absent from the 114 HHC individuals. 1,236 convergent clusters met these criteria and showed SHM frequencies averaging 1.7% (range 0.5% to 5.5%). An average of 196 convergent clusters were found per patient, ranging from 69 clusters in patient 7485 to 477 clusters in patient 7455. 1,171 clusters were shared pairwise between two patients, 53 clusters spanned three patients, nine clusters spanned four patients, and three clusters spanned five patients (Figure 3A). To assess the significance of these shared convergent clones in COVID-19 patients, we undertook an analysis of 13 randomly selected HHCs with the same parameters and 100 permutations. The number of convergent clones shared by COVID-19 patients greatly exceeded the mean convergent clone counts from the HHC subsampling (Figure 3B), consistent with antigen-driven shared selection of the convergent clones identified in COVID-19 patients. To directly test the antigen specificity of these convergent clones, we expressed human IgG1 monoclonal antibodies (mAbs 2A and 4A, Table S2) belonging to two COVID-19 convergent antibody groups identified in two patients from whom paired immunoglobulin heavy and light chain sequences were obtained from single B cells using the 10x Genomics platform. mAb2A had a single nonsynonymous mutation in the CDR-H2 of IGHV3-30-3 and mAb4A was fully germline and used IGHV3-15. In ELISA testing, both mAbs bound the SARS-CoV-2 spike protein and spike S1 domain, but not the RBD (Figure 3C), establishing their antigen specificity. We further validated the robustness of detection of convergent clonotypes in the COVID-19 patients, and the wide distribution of these antibody types in different individuals, by identifying ten SARS-CoV-2 spike RBD-specific sequence clusters shared with independent external COVID-19 patient data sets (Cao et al., 2020; Noy-Porat et al., 2020; Shi et al., 2020). Nine of the thirteen COVID-19 patients (69%) in our study (7450, 7452, 7454, 7455, 7480, 7483, 7484, 7485, and 7486) showed these RBD-specific clonotypes, highlighting the high frequency of these shared antibodies (Figure 3D and Figure S4). Notably, RBD-specific antibodies are primary candidates for virus neutralizing, potentially protective antibodies in recovered patients (Ju et al., 2020; Robbiani et al., 2020a).

**Figure 3.**
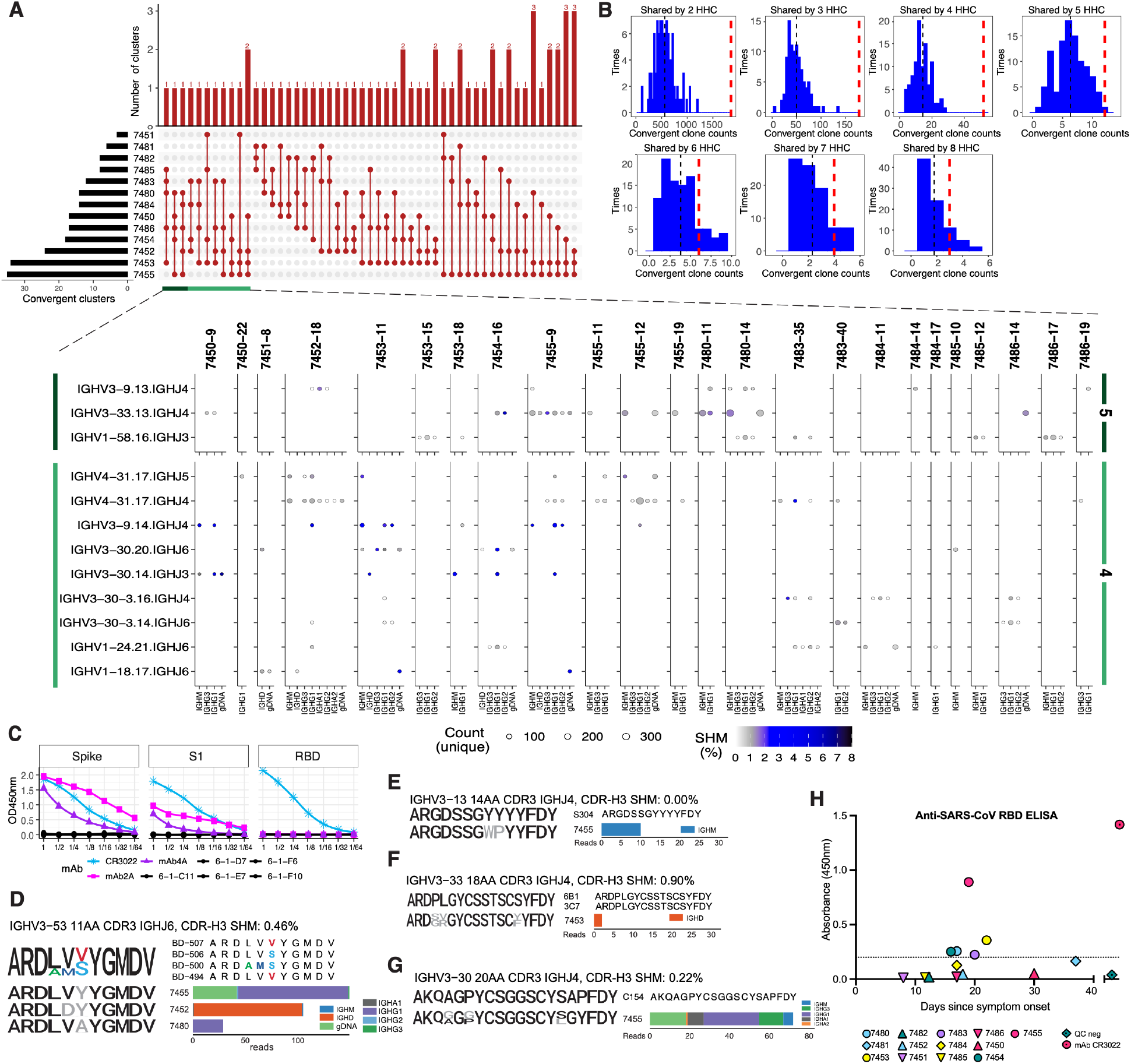
Convergent IGH sequences between COVID-19 patients and to reported antigen-specific IGH correlate with SARS-CoV/SARS-CoV-2 plasma cross-reactivity. (A) The distribution of convergent clusters among patient samples (top panel). The sample distribution is indicated by the lines and dots with the number of clusters sharing that sample distribution indicated by the vertical histogram bars. The total number of convergent clusters identified in each sample is indicated in the histogram to the left of the plot. (bottom panel) Lineages belonging to convergent clusters (y-axis) shared across four or five patients (columns, four-patient clusters are highlighted in light green, five-patient clusters in dark green) are plotted by the expressed isotype (x-axis). Fill color indicates the average SHM and point size shows the number of unique reads. (B) Distribution of the number of convergent clones identified in two to eight HHC subjects, determined using 100 permutations. The histogram shows the distribution of convergent clones shared by a given number of HHC samples each time. The black dashed line is the mean value, and the wide red dashed line is the number of convergent clones shared among the same number of COVID-19 participants. (C) ELISA assay results for human IgG1 mAb binding to SARS-CoV-2 spike ectodomain protein, spike S1 domain, or RBD. Negative control mAbs 6-1-C11,D7,E7,F6, and F10 are overplotted in black. The SARS-CoV-2 RBD-binding mAb CR3022 (ter Meulen et al., 2006; Tian et al., 2020) was used as positive control. Starting concentration for mAbs was 100 ug/mL save for CR3022, which started at 0.506 ug/mL. (D-G) Sequence logos of CDR-H3 AA residues from anti-SARS-CoV-2 (D, see also Figure S4) anti-SARS-CoV/CoV-2 cross-neutralizing (E) or anti-SARS-CoV (F-G) convergent IGHs. For each set of convergent IGH the sequence logo and alignment for the reported antigen-specific CDR-H3 is shown at the top, sequence logos for clones from each patient are aligned below (colored black where they match a conserved residue in the reported CDR-H3, colored for non-conserved as depicted in the alignment, or gray if no match). To the side the read count per patient that contributed to the sequence logo, by isotype, is graphed. The SHM frequency for the dominant isotype is shown after the convergent IGH label. (H) Anti-SARS-CoV IgG ELISA detection in plasma samples from COVID-19 patients. Plasma samples were analyzed for the presence of anti-SARS-CoV spike RBD-binding IgG antibodies in the latest sample timepoint available for each patient. A SARS-CoV-2 pre-pandemic sample pool from healthy blood donors was used as a negative quality control (QC) as well as a positive control for SARS-CoV RBD (mAb CR3022) (ter Meulen et al., 2006). The dotted line denotes the cut-off value for seroconversion. Assays were performed in duplicate and mean OD values are shown.

We hypothesized that in addition to sharing common RBD-binding antibody types, some COVID-19 patients might also demonstrate breadth in their antibody responses and recognize antigens from the distinct but related sarbecovirus, SARS-CoV that was responsible for SARS. Comparison of COVID-19 patient IGH sequences to published SARS patient IGH data revealed convergent SARS-CoV RBD-specific antibodies in two COVID-19 patients, 7453 and 7455 (Coughlin et al., 2007; Pinto et al., 2020; Robbiani et al., 2020a) (Figures 3E-3G). None of these SARS-CoV-specific convergent IGH sequences were detected in the 114 HHC. To evaluate whether such *in silico* IGH sequence comparisons could predict the serological responses of patients, we tested the plasma samples from our 13 COVID-19 patients in ELISA assays with SARS-CoV RBD antigen and detected cross-reactivity in five of the 13 patients (Figure 3H). Strikingly, the two patients with the highest ELISA OD450 values were those who had demonstrated convergent IGH sequences specific for SARS-CoV RBD. The three additional COVID-19 patients who were seropositive for SARS-CoV RBD antibodies had convergent IGH sequences to SARS-CoV-2 in their repertoires, suggesting that the presence of these convergent antibodies could be a marker of more extensive or broadly-reactive humoral immune responses in patients.

## DISCUSSION

In these initial months of the COVID-19 pandemic, understanding human antibody responses to SARS-CoV-2 has become a global priority. Our results provide several key findings that may lend some support for vaccine strategies currently under development and suggest that individuals convalescent from SARS-CoV-2 infection may be, at least for some time, protected against reinfection by commonly-elicited RBD-specific antibodies. The IGH repertoires of patients with diagnostically confirmed SARS-CoV-2 reveal robust polyclonal responses with early class switching to IgG, and to a lesser extent, IgA isotypes, and evidence of accumulating SHM in responding clones within the first month after onset of symptoms, rather than the delayed SHM seen in Ebola patients (Davis et al., 2019). We note that the current COVID-19 study and prior analysis of EBOV infection are among very few published studies of human IGH repertoire longitudinal responses to primary infections; examples from acute infection with Dengue virus (Appanna et al., 2016; Godoy-Lozano et al., 2016) or H5N6 avian influenza virus (Peng et al., 2019), have either had few patients with true primary infection, or did not analyze SHM development in responding B cells.

Nine of thirteen COVID-19 patients (69%) demonstrated convergent antibodies specific for the viral RBD, a major target for potentially neutralizing antibodies. SARS-CoV-2 neutralizing serum antibodies are reported to be present in 67-90% of patients post-infection, depending on the severity of disease, neutralization assay and threshold for positive results (Robbiani et al., 2020b; Suthar et al., 2020; Wu et al., 2020a). It seems reasonable to predict that vaccines based on spike or RBD antigens will also stimulate B cells expressing these common antibody types in a significant fraction of the human population. The response to SARS-CoV-2 infection in a subset of patients also contained B cell clones expressing convergent IGH to previously described SARS-CoV RBD antibodies; strikingly, the patients with these SARS-CoV-2/SARS-CoV clonotypes also had the highest SARS-CoV RBD binding serum antibody IgG levels. This association suggests that it may become possible to predict the fine specificity of human serological responses from IGH sequence data, as the number of documented antigen-specific clonotypes in public databases increases. This example also highlights the possibility that common modes of human antibody response may enable some breadth of protection or humoral memory against other sarbecoviruses in the future. Longitudinal tracking of IGH repertoires in larger patient cohorts, further investigation into the binding properties, functional activity and serum antibody levels produced by convergent responding clones in patients, and assessment of clinical outcomes under conditions of exposure to infection will be important next steps toward determining the immunological correlates of protection against SARS-CoV-2 infection.

## ACKNOWLEDGMENTS

We thank the patients, Hannah K. Frank for initial advice on sequence alignments, Shilpa A. Joshi for editing the paper, Jonasel Roque, Philip Grant, Aruna Subramanian for assistance with recruiting and consenting subjects, Aaron J. Wilk, Nancy Q. Zhao, Giovanny J. Martinez-Colon, Julia L. McKechnie, Geoffrey Ivison, Thanmayi Ranganath, Rosemary Vergara, and Laura J. Simpson for processing of samples.

## AUTHOR CONTRIBUTIONS

C.A.B. and S.D.B. conceived the project. A.R., A.J.R. recruited, enrolled, and consented patients and contributed to clinical sampling and processing. S.C.A.N., R.A.H., K.R., B.S., J-Y.L., A.E.P., J.N., A.R.O-C., K.E.Y., B.D. and B.A.P. performed the experiments. H.Y.C., A.T.S., T.S.J., P.S.K. and T.T.W. contributed to serological or 10x assays, and analysis. S.C.A.N., F.Y., R.A.H., K.J.L.J., K.R., and B.A.P. contributed to data analysis. S.C.A.N., F.Y., R.A.H., K.J.L.J., K.R., C.A.B., and S.D.B. wrote the manuscript. All authors edited the manuscript.

## CONTACT FOR REAGENT AND RESOURCE SHARING

Further information and requests for resources and reagents should be directed to the Lead Contact, Scott D. Boyd.

## DATA AND SOFTWARE AVAILABILITY

All data is available in the main text or the extended materials. The IGH repertoire data for this study have been deposited to SRA with accession number PRJNA628125.

## EXPERIMENTAL MODELS AND SUBJECT DETAILS

Patients admitted to Stanford Hospital with signs and symptoms of COVID-19 and confirmed SARS-CoV-2 infection by RT-qPCR of nasopharyngeal swabs were recruited. Venipuncture blood samples were collected in K_2_EDTA- or sodium heparin-coated vacutainers for peripheral blood mononuclear cell (PBMC) isolation or serology on plasma, respectively. Recruitment of COVID-19 patients, documentation of informed consent, collections of blood samples, and experimental measurements were carried out with Institutional Review Board approval (IRB-55689). The data set containing healthy adult control immunoglobulin receptor repertoires has been described previously (Nielsen et al., 2019). In summary, healthy adults with no signs or symptoms of acute illness or disease were recruited as volunteer blood donors at the Stanford Blood Center. Pathogen diagnostics were performed for CMV, HIV, HCV, HBV, West Nile virus, HTLV, TPPA (Syphilis), and *T. cruzi*. Volunteer age range was 17-87 with median and mean of 52 and 49, respectively.

## METHOD DETAILS

### Molecular and serological testing on COVID-19 patient samples

SARS-CoV-2 infection in patients was confirmed by reverse-transcription polymerase chain reaction testing of nasopharyngeal swab specimens, using the protocols described in (Corman et al., 2020; Hogan et al., 2020). An enzyme-linked immunosorbent assay (ELISA) based on a protocol described in (Stadlbauer et al., 2020) was performed to detect anti-SARS-CoV and anti-SARS-CoV-2 spike RBD antibodies in plasma samples from COVID-19 patients. Briefly, 96-well high binding plates (Thermo Fisher) were coated with either SARS-CoV or SARS-CoV-2 spike RBD protein (0.1 μg per well) overnight at 4°C. After blocking plates with 3% non-fat milk in PBS containing 0.1% Tween 20, plasma samples were incubated at a dilution of 1:100 and bound antibodies were detected with goat anti-human IgM/HRP (Sigma: cat. A6907, 1:6’000 dilution), goat anti-human IgG/HRP (Thermo Fisher: cat. 62-8420, 1:6’000 dilution), or rabbit anti-human IgA/HRP (Dako: cat. P0216, 1:5’000 dilution). Assays were developed by addition of 3,3’,5,5’-Tetramethylbenzidine (TMB) substrate solution. After stopping the reaction with 0.16 M sulfuric acid, the optical density (OD) at 450 nanometers was read using an EMax Plus microplate reader (Molecular Devices). The cut-off value for seroconversion was calculated as OD_450_ = 0.2 for the anti-SARS-CoV IgG assay and as OD_450_ = 0.3 for anti-SARS-CoV-2 IgM, IgG, and IgA assays after analyzing SARS-CoV-2 pre-pandemic negative control samples from healthy blood donors.

### HTS of immunoglobulin heavy chain (IGH) libraries prepared from genomic DNA and cDNA

The AllPrep DNA/RNA kit (Qiagen) was used to extract genomic DNA (gDNA) and total RNA from PBMCs. For each blood sample, six independent gDNA library PCRs were set up using 100 ng template/library (25ng/library for 7453-D0). Multiplexed primers to IGHJ and the FR1 or FR2 framework regions (3 FR1 and 3 FR2 libraries), per the BIOMED-2 design were used (van Dongen et al., 2003) with additional sequence representing the first part of the Illumina linkers. In addition, for each sample, total RNA was reverse-transcribed to cDNA using Superscript III RT (Invitrogen) with random hexamer primers (Promega). Total RNA yield varied between patients and between 6 ng-100 ng was used for each of the isotype PCRs using IGHV FR1 primers based on the BIOMED-2 design (van Dongen et al., 2003) and isotype specific primers located in the first exon of the constant region for each isotype category (IgM, IgD, IgE, IgA, IgG). Primers contain additional sequence representing the first part of the Illumina linkers. The different isotypes were amplified in separate reaction tubes. Eight-nucleotide barcode sequences were included in the primers to indicate sample (isotype and gDNA libraries) and replicate identity (gDNA libraries). Four randomized bases were included upstream of the barcodes on the IGHJ primer (gDNA libraries) and constant region primer (isotype libraries) for Illumina clustering. PCR was carried out with AmpliTaq Gold (Applied Biosystems) following the manufacturer’s instructions, and used a program of: 95°C 7 min; 35 cycles of 94°C 30 sec, 58°C 45 sec, 72°C 60 sec; and final extension at 72°C for 10 min. A second round of PCR using Qiagen’s Multiplex PCR Kit was performed to complete the Illumina sequencing adapters at the 5’ and 3’ ends of amplicons; cycling conditions were: 95°C 15 min; 12 cycles of 95°C 30 sec, 60°C 45 sec, 72°C 60 sec; and final extension at 72°C for 10 min. Products were subsequently pooled, gel purified (Qiagen), and quantified with the Qubit fluorometer (Invitrogen). Samples were sequenced on the Illumina MiSeq (PE300) using 600 cycle kits.

### Sequence quality assessment, filtering, and analysis

Paired-end reads were merged using FLASH (Magoc and Salzberg, 2011), demultiplexed (100% barcode match), and primer trimmed. The V, D, and J gene segments and V-D (N1), and D-J (N2) junctions were identified using the IgBLAST alignment program (Ye et al., 2013). Quality filtering of sequences included keeping only productive reads with a CDR-H3 region, and minimum V-gene alignment score of 200. Sample cDNA or gDNA libraries with poor read coverage were excluded from further analysis (Table 1). For cDNA-templated IGH reads, isotypes and subclasses were called by exact matching to the constant region gene sequence upstream from the primer. Clonal identities within each subject were inferred using single-linkage clustering and the following definition: same IGHV and IGHJ usage (disregarding allele call), equal CDR-H3 length, and minimum 90% CDR-H3 nucleotide identity. A total of 1,259,882 clones (per sample, mean number of clones: 33,154; median number of clones: 18,503) were identified. A total of 24,888,790 IGH sequences amplified from cDNA were analyzed for the COVID-19 subjects (mean: 754,205 per sample; median: 650,812) and 68,831,446 sequences from healthy adult controls (mean: 603,785 per individual; median: 637,269). Each COVID-19 patient had on average 372,304 in-frame gDNA sequences and each adult control had an average of 8,402 in-frame gDNA sequences.

For each clone, the median somatic mutation frequency of reads was calculated. Mean mutation frequencies for all clonal lineages from a sample for each isotype were calculated from the median mutation frequency within each clone, and so represent the mean of the median values. Clones with <1% mutation were defined as unmutated and clones with ≥ 1% were defined as being mutated. Subclass fractions were determined for each subject by dividing the number of clones for a given subclass by the total number of clones for that isotype category. Expanded clones within each sample were defined as clones that were present in two or more of the gDNA replicate libraries. Clonal expansion in the isotype data was inferred from the gDNA data. Analyses were conducted in R (Team, 2017) using base packages for statistical analysis and the ggplot2 package for graphics (Wickham, 2016).

To determine convergent rearranged IGH among patients with SARS-CoV-2 infection, we clustered heavy-chain sequences annotated with the same IGHV and IGHJ segment (not considering alleles) and the same CDR-H3 length were clustered based on 85% CDR-H3 amino acid sequence similarity using CD-HIT (Fu et al., 2012). To exclude IGH that are generally shared between humans and to enrich the SARS-CoV-2-specific IGH that are likely shared among the patients, clusters were selected as informative if (1) they contained at least five IGH sequences from each COVID-19 patient and were present in at least two subjects; (2) no IGH sequences from HHC samples (collected prior to the 2019 SARS-CoV-2 outbreak) were identified in the same convergent cluster. The same selection criteria were used to determine the convergent clusters between the COVID-19 samples and previously reported IGH sequences specific to SARS-CoV and SARS-CoV-2. Convergent IGH sequences between the deeply sequenced COVID-19 patients and 10x Genomics single B cell immune profiling on two COVID-19 patients were selected for mAb expression.

### Single-cell immunoglobulin (Ig) library preparation, sequencing, and data processing

Single-cell immunoglobulin libraries were prepared using the 10x Single Cell Immune Profiling Solution Kit (v1.1 Chemistry), according to the manufacturer’s instructions. Briefly, cells were washed once with PBS + 0.04% BSA. Following reverse transcription and cell barcoding in droplets, emulsions were broken and cDNA purified using Dynabeads MyOne SILANE followed by PCR amplification (98°C for 45 sec; 15 cycles of 98°C for 20 sec, 67°C for 30 sec, 72°C for 1 min; 72°C for 1 min). For targeted Ig library construction, 2 μL of amplified cDNA was used for target enrichment by PCR (Human B cell primer sets 1 and 2: 98°C for 45 sec; 6 and 8 cycles of 98°C for 20 sec, 67°C for 30 sec, 72°C for 1 min; 72°C for 1 min). Following Ig enrichment, up to 50 ng of enriched PCR product was fragmented and end-repaired, size selected with SPRIselect beads, PCR amplified with sample indexing primers (98°C for 45 sec; 9 cycles of 98°C for 20 sec, 54°C for 30 sec, 72°C for 20 sec; 72°C for 1 min), and size selected with SPRIselect beads. Targeted single-cell Ig libraries were sequenced on an Illumina MiSeq to a minimum sequencing depth of 5,000 reads/cell using the read lengths 26bp Read1, 8bp i7 Index, 91bp Read2 and reads were aligned to the GRCh38 reference genome and consensus Ig annotation was performed using cellranger vdj (10x Genomics, version 3.1.0).

### Identification of convergent IGH sequences for mAb expression

IGH sequences from single cells with paired productive heavy and light chains were searched against COVID-19 patient bulk IGH repertoires to identify convergent sequences according to the following criteria: utilization of the same IGHV and IGHJ genes; same CDR-H3 lengths; and CDR-H3 amino acid sequences that were within a Hamming distance cutoff of 15% of the length of the CDR-H3. Two native heavy and light chain pairs, designated mAb2A and mAb4A, which were found in convergent clusters characterized by low-to mid-SHM frequencies and included at least one class-switched member, were selected for cloning and expression.

### mAb cloning

Paired heavy and light chain sequences from 10x single cell RNA-seq datasets were synthesized by IDT as gBlocks encoding full-length heavy and light chain V(D)J regions. gBlocks were resuspended at 50 ng/μL and amplified with AmpliTaq Gold (Applied Biosystems) following the manufacturer’s instructions, using a program of: 95°C 7 min; 30 cycles of 94°C for 30 sec, 55°C for 45 sec, 72°C for 60 sec; and final extension at 72°C for 10 min. Products were gel purified (Qiagen) and cloned as in-frame fusions to human IgG1, IgK or IgL constant regions into the pYD7 vector (National Research Council (NRC), Canada) using Gibson Assembly Master Mix (NEB) for 45 minutes at 50°C. Assembled constructs were verified by Sanger sequencing.

### mAb expression and purification

Constructs were transiently transfected in HEK293-EBNA1-6E cells (NRC) at a density of 1.2-1.6 million cells/mL using 25 kDa linear polyethylenimine (PEI) at a 3:1 PEI:DNA ratio in OptiMEM reduced serum medium (Gibco), with a heavy chain: light chain ratio of 1:1. Cells were maintained in Freestyle 293 Expression Medium (Gibco) and were supplemented with 0.5% tryptone 24-36 hours after transfection. Cell supernatants were harvested after 96 hours and filtered through 0.45-μM filters (Millipore). Antibodies were purified via HiTrap Protein A HP columns (GE Healthcare) run at a flow rate of 0.5-1 mL/min on Äkta Start protein purification system (GE Healthcare). Antibodies were eluted using 0.1M glycine pH 2, dialyzed with 3 changes of PBS pH 7.4 using Slide-A-Lyzer-G2 10K dialysis cassettes (Thermo Fisher), and concentrated using 30,000 kDa molecular weight cutoff polyethersulfone membrane spin columns (Pierce). Final concentrations of purified antibodies were quantified with Nanodrop 2000 (Thermo Fisher).

### ELISA testing of mAbs

ELISA conditions for mAbs were as described for COVID-19 plasma samples with the following modifications: two-fold serial dilutions of mAbs were tested, starting at 100 μg/mL for intra-COVID-19 convergent antibodies or peanut-specific negative mAb controls, or at 0.506 μg/mL for mAb CR3022; plates were coated overnight with RBD (0.1 μg per well), S1 (0.1 μg per well), or spike protein (0.3 μg per well); and bound mAbs were detected with rabbit anti-human IgG gamma chain-specific/HRP (Agilent: cat. P0214, 1:15,000 dilution).

## SUPPLEMENTAL INFORMATION

Supplemental Information includes two tables and four figures.

