## Supplemental File for "Human B cell clonal expansion and convergent antibody responses to SARS-CoV-2"

### Supplemental Information

**Table S1, related to Figure 1. IgG and IgA subclass proportions.** COVID-19 and healthy human control (HHC) median isotype subclass proportions +/- median absolute deviation (MAD) summarized for all samples within each group. p-values were calculated by two-sided Wilcoxon–Mann-Whitney tests.

| | **COVID-19** (median +/- MAD) | **HHC** (median +/- MAD) | **Two-sided Wilcoxon–Mann-Whitney test** |
|---|---|---|---|
| IGHA1 | 70.1 +/- 9.3 | 65 +/- 6.6 | p-value = 0.2018 |
| IGHA2 | 29.9 +/- 9.4 | 35 +/- 6.6 | p-value = 0.2018 |
| IGHG1 | 69.2 +/- 11.5 | 41.4 +/- 7.3 | p-value = 9.114e-09 |
| IGHG2 | 16.3 +/- 11.2 | 40.2 +/- 8.5 | p-value = 8.855e-08 |
| IGHG3 | 8.7 +/- 3.3 | 14.6 +/- 5.9 | p-value = 6.281e-05 |
| IGHG4 | 0.3 +/- 0.2 | 1.6 +/- 1.9 | p-value = 0.003414 |

**Table S2, related to Figure 3. Heavy and light chain sequence features for convergent monoclonal antibodies (mAbs) 2A and 4A.** The percent identity (ID %) of the IGHV, IGKV or IGLV gene sequence relative to germline is shown.

| **mAb** | **IGHV** | **IGHD** | **IGHJ** | **IGHC** | **IGHV ID (%)** | **CDR-H3 AA** | **IGK/LV** | **IGK/LJ** | **IGK/LV ID (%)** | **CDR-L3 AA** |
|---|---|---|---|---|---|---|---|---|---|---|
| **mAb2A** | IGHV3-30-3*01 | IGHD3-22*01 | IGHJ3*02 | IGHG1 | 99.7 | ARDSGSAFDI | IGLV3-1*01 | IGLJ2*01, IGLJ3*01 | 100 | QAWDSSTVV |
| **mAb4A** | IGHV3-15*01 | IGHD3-16*02 | IGHJ4*02 | IGHM | 100 | TTDRHYDYVWGSYRYPDY | IGKV1-39*01 | IGKJ5*01 | 100 | QQSYSTPT |

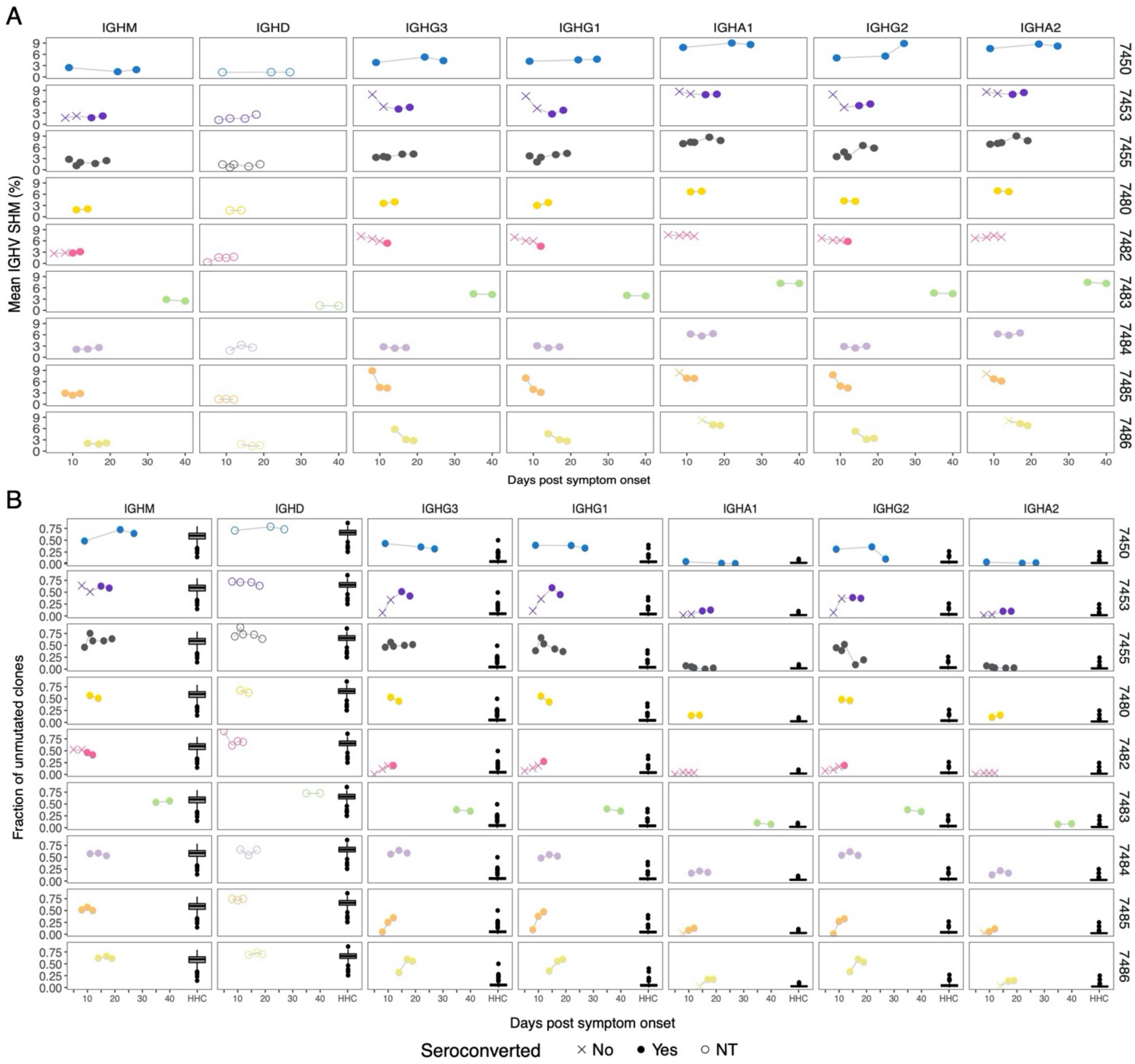


**Figure S1, related to Figure 1. Mean SHM percentage and unmutated clone fractions over time for each patient. (A)** The average SHM (%) for the IGHV gene segment of expressed antibodies of the indicated isotype (panel column) for each individual (panel row). SHM for each isotype in each sample was summarized as the median SHM of reads expressed as the indicated isotype within each clone, then taking the mean SHM percentage over all clones. **(B)** The fraction of unmutated (<1% IGHV SHM, y-axis) B cell lineages for each IGH of the indicated isotype (panel column) for each individual (panel row). **(A)** and **(B)** Days post symptom onset on x-axis. Point shapes indicate patient sample serology; seronegative (x), and seropositive (filled circle), not tested (NT, open circle) and are plotted specific to the isotype tested.

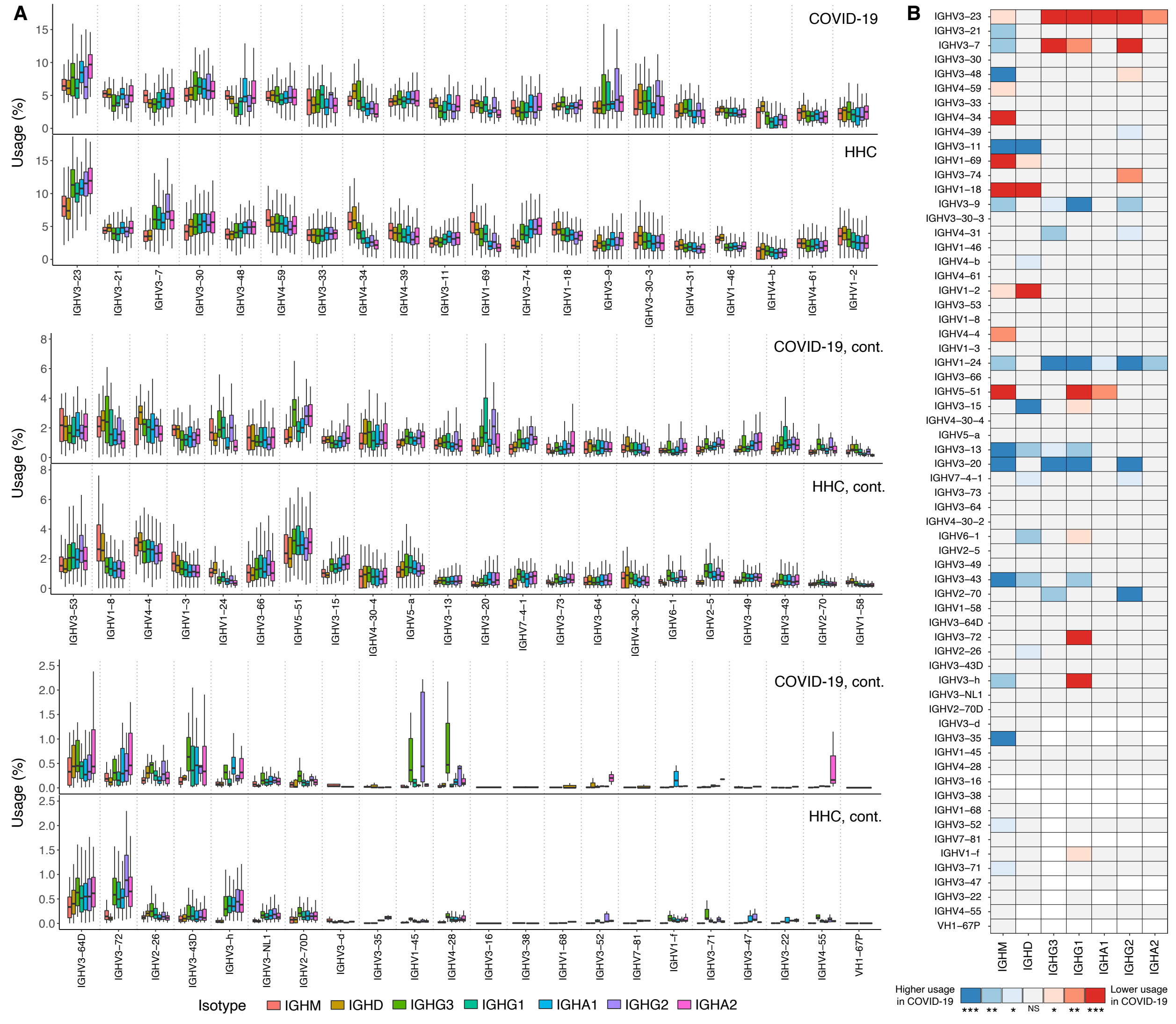


**Figure S2, related to Figure 2. IGHV usage for COVID-19 patients compared to a healthy human control (HHC) cohort.** Median frequency of IGHV gene utilization for each isotype subclass observed for the 13 COVID-19 patients compared to a cohort of 114 HHCs for all IGHV genes utilized. **(A)** IGHV usage is shown as the median for clones within each COVID-19 patient and HHC. **(B)** Gene utilization frequencies for each isotype subclass for each IGHV gene were compared between COVID-19 and HHC (paired Wilcoxon tests with Bonferroni correction for multiple hypothesis testing). Where gene usage differed significantly between the cohorts and the gene was utilized at a higher median frequency among the COVID-19 patients the adjusted significance is shown in blue, whereas genes that differed significantly but with lower median usage for the COVID-19 patients relative to the HHC are plotted in red. Instances with insufficient data for test (only 1 or no data points) are in white. The plot y-axes were chosen to show the box-whiskers on a readable scale; rare outlier points with extreme values are not shown but were included in all analyses.

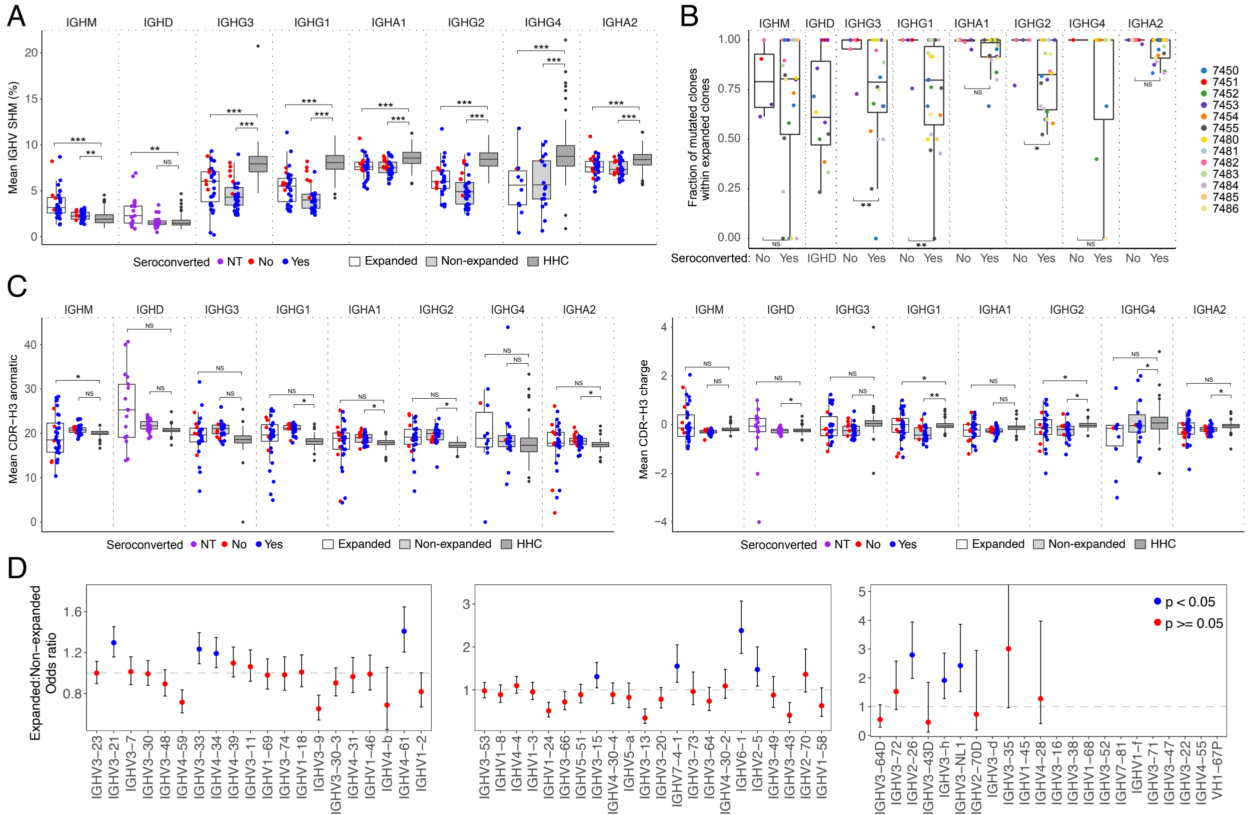


**Figure S3, related to Figure 2. IGHV gene and CDR-H3 features between expanded and non-expanded patient clones and healthy human controls (HHC). (A)** Mean IGHV SHM of expanded and non-expanded clones in COVID-19 patients compared to HHC. SHM frequency for each isotype in each sample was summarized as the median SHM of reads expressed as the indicated isotype within each clone, then taking the mean SHM percentage over all clones. Points are jittered on the x-axis to decrease over-plotting of samples with the same value (y-axis). p-values were calculated by two-sided Wilcoxon–Mann-Whitney tests. **(A)** and **(C)** COVID-19 patient samples grouped by expanded clones (white) or non-expanded clones (light grey), and total clones from healthy human (HHC) (dark grey, only outlier points are displayed for this group). Points are colored by seroconversion status; not tested (NT, purple), seronegative (red), and seropositive (blue) and are plotted specific to the isotype tested. **(B)** Fraction of mutated expanded clones (y-axis) by seroconversion status (x-axis). Points are colored by participant. p-values were calculated by Fisher's exact tests. **(C)** Mean CDR-H3 charge and aromaticity of expanded clones in COVID-19 patients. Percent aromaticity (left panel) and mean charge (right panel) for CDR-H3 amino acid residues. p-values were calculated by one-way ANOVA with Tukey's HSD test. **(A-C)**, ***p-value < 0.001; **p-value < 0.01; *p-value ≤ 0.05; NS: p-value > 0.05. **(D)** Odds ratio of IGHV gene usage in expanded clones compared to non-expanded clones in COVID-19 patients. Each dot is the odds ratio (OR) value for each IGHV gene, the bars represent confidence interval. The plot y-axis was chosen to show the points on a readable scale; extreme values are not shown but were included in all analyses. Each point was colored by p-value tested by Fisher's exact test, blue: p-value < 0.05, red: p-value >= 0.05.

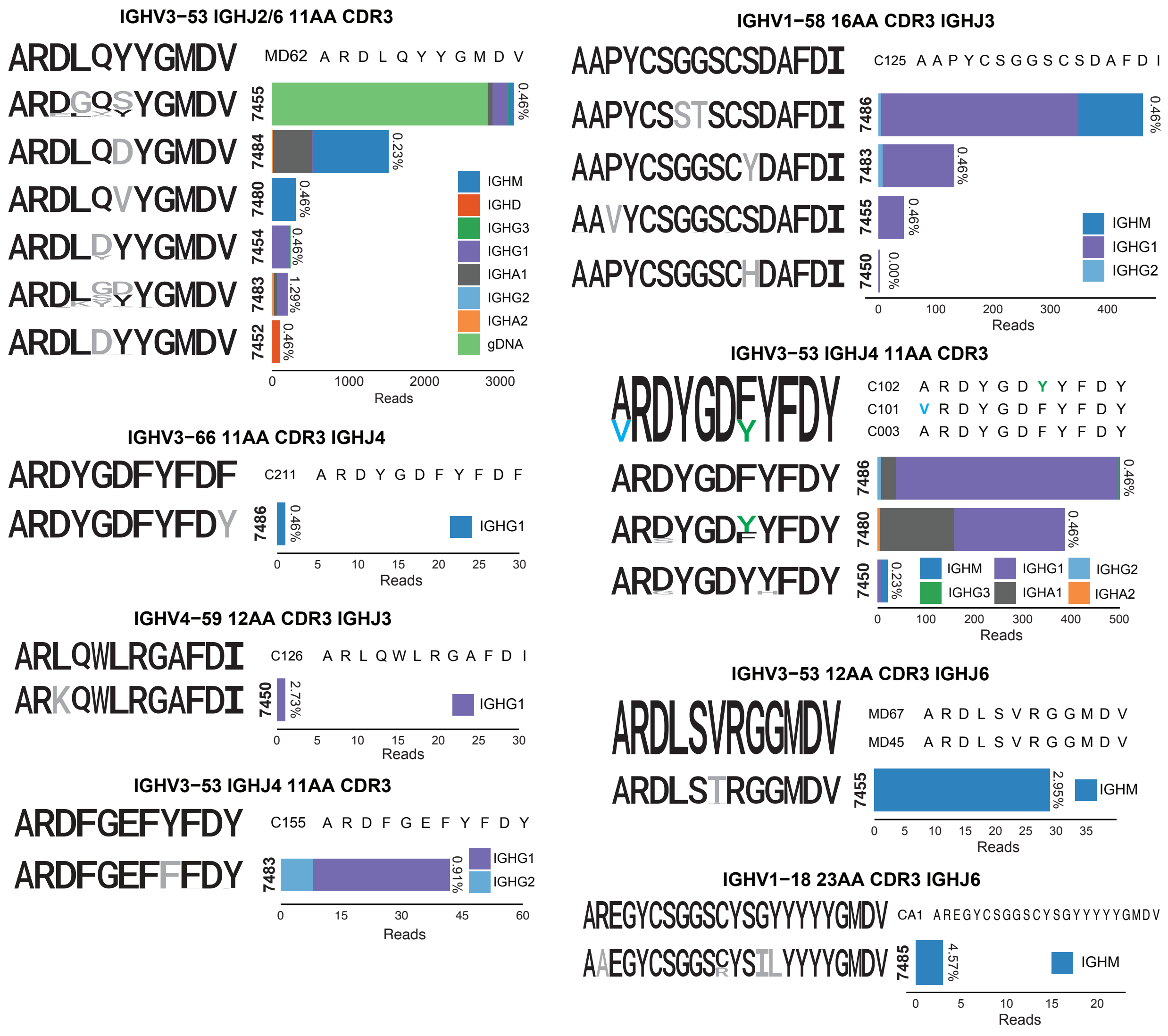


**Figure S4, related to Figure 3. Sequence logos of CDR-H3 AA residues from anti-SARS-CoV-2 convergent IGHs.** For each set of convergent IGH the sequence logo and alignment for the reported antigen-specific CDR-H3 is shown at the top, and sequence logos for clones from each patient are aligned below (colored black where they match a conserved residue in the reported CDR-H3, colored for non-conserved as depicted in the alignment, or gray if no match). To the side the read count per patient that contributed to the sequence logo, by isotype, is graphed with the SHM frequency printed above the bar.
